# Susceptibility of well-differentiated airway epithelial cell cultures from domestic and wildlife animals to SARS-CoV-2

**DOI:** 10.1101/2020.11.10.374587

**Authors:** Mitra Gultom, Matthias Licheri, Laura Laloli, Manon Wider, Marina Strässle, Silvio Steiner, Annika Kratzel, Tran Thi Nhu Thao, Hanspeter Stalder, Jasmine Portmann, Melle Holwerda, Philip V’kovski, Nadine Ebert, Nadine Stokar – Regenscheit, Corinne Gurtner, Patrik Zanolari, Horst Posthaus, Simone Schuller, Amanda Vicente – Santos, Andres Moreira – Soto, Eugenia Corrales – Aguilar, Nicolas Ruggli, Gergely Tekes, Veronika von Messling, Bevan Sawatsky, Volker Thiel, Ronald Dijkman

**Author notes:** Current address: Institute for Infectious Diseases, University of Bern, Bern, Switzerland (Philip V’kovski). corresponding author: Ronald Dijkman.

## Abstract

Severe Acute Respiratory Syndrome Coronavirus 2 (SARS-CoV-2) has spread globally, and the number of cases continues to rise all over the world. Besides humans, the zoonotic origin, as well as intermediate and potential spillback host reservoirs of SARS-CoV-2 are unknown. To circumvent ethical and experimental constraints, and more importantly, to reduce and refine animal experimentation, we employed our airway epithelial cell (AEC) culture repository composed of various domesticated and wildlife animal species to assess their susceptibility to SARS-CoV-2. In this study, we inoculated well-differentiated animal AEC cultures of monkey, cat, ferret, dog, rabbit, pig, cattle, goat, llama, camel, and two neotropical bat species with SARS-CoV-2. We observed that SARS-CoV-2 only replicated efficiently in monkey and cat AEC culture models. Whole-genome sequencing of progeny virus revealed no obvious signs of nucleotide transitions required for SARS-CoV-2 to productively infect monkey and cat epithelial airway cells. Our findings, together with the previously reported human-to-animal spillover events warrants close surveillance to understand the potential role of cats, monkeys, and closely related species as spillback reservoirs for SARS-CoV-2.

## Introduction

During the last two decades we have observed zoonotic outbreaks of Severe Acute Respiratory Syndrome Coronavirus (SARS-CoV) in 2003 and Middle East Respiratory Syndrome Coronavirus (MERS-CoV) in 2012 ^1,2^. These outbreaks are followed by the current pandemic which caused by zoonotic emergence of SARS-CoV-2, the etiological agent of Coronavirus Disease 2019 (COVID-19) ^3,4^. Although humans are currently seen as the main hosts, the zoonotic origin, as well as the intermediate and potential spillback host reservoirs of SARS-CoV-2 is unknown. Interestingly, several reports indicate that SARS-CoV-2 spillover events from human to other animal species can occur ^5–9^. These events are likely driven by close human-animal interactions and the conservation of crucial receptor binding motif (RBM) residues in the angiotensin-converting enzyme 2 (ACE2) orthologues, potentially facilitating SARS-CoV-2 entry ^10,11^. This highlights the need to assess the potential host spectrum for SARS-CoV-2 in order to support current pandemic mitigation strategies.

Besides risk assessment of the host spectrum, viral pathogenesis studies, as well as all novel antiviral drugs, immunotherapies, and vaccines against SARS-CoV-2, will require evaluation in animal models. Therefore, a large variety of animal species are tested on their susceptibility ^12–14^. Traditionally such experiments have several drawbacks in terms of animal model diversity and availability, including dedicated personnel, housing facilities, and most importantly, ethical approval. Some of these factors are especially limited in the context of wildlife, and livestock animals, such as pig, cattle and other ruminants, and in case of companion animals and non-human primates there are additional socioemotional ethical constraints to be overcome.

In this study, we evaluated the susceptibility of several mammalian species to SARS-CoV-2 by recapitulating the initial stages of infection in a controlled *in vitro* model, in compliance with the reduction, refinement and replacement principles in animal experimentation, whilst at the same time circumventing traditional *in vivo* experimental constraints. We employed a unique well-differentiated airway epithelial cell (AEC) culture repository from primary tracheobronchial airway tissue of 12 mammalian species comprising companion animals, animal model candidates, livestock, and wild animals to assess their susceptibility to SARS-CoV-2 infection. In order to control for the quality of the AEC, we used influenza viruses that have a known broad host tropism ^15–17^.

## Results

The main goal of our study is to evaluate the susceptibility of a diverse set of animal species to SARS-CoV-2 infection. We employed well-differentiated AEC cultures reconstituted from tracheobronchial epithelial tissue, thereby replacing animal experimentation. In total, we obtained post-mortem tracheobronchial airway tissue material from animal species, comprising companion animals (cat, dog), commonly used animal models (rhesus macaque, ferret, and rabbit), livestock (pig, cattle, goat, camel, and llama), and two neotropical frugivorous bat species (*Sturnira lilium* and *Carollia perspicillata*). Following isolation of primary airway epithelial cells, well-differentiated AEC cultures were established and maintained as performed previously for human AEC cultures, with individual modifications for each species (**Table. 1**) ^18^. After 4 to 6 weeks of differentiation we inoculated each culture with 30.000 PFU of the SARS-CoV-2/München-1.1/2020/929 isolate, representing the current circulating Spike^D614G^ variant 19. Viral progeny release was monitored by quantifying the viral titer and viral RNA in the apical washes that were collected every 24 hours for a duration of 96 hours. Since we previously observed that SARS-CoV-2 replication efficiency in the human respiratory tract is influenced by the ambient temperature, the infection was carried out at either 33°C or 37°C ^20^.

**Table 1.** Optimized epidermal growth factor (EGF), retinoic acid (RA), hydrocortisone (HC), and DAPT concentration in the ALI medium for differentiation of the animal AEC cultures.

| No | Animal species | Number of donors | Source | End concentration additives in ALI medium |  |  |  |
| --- | --- | --- | --- | --- | --- | --- | --- |
|  |  |  |  | EGF | RA | HC | DAPT |
| 1 | Monkey<br>(*Rhesus macaque*) | 2 | Paul-Ehrlich-Institut, Langen, Germany | 5 ng/ml | 50 nM | 0.48 µg/ml | - |
| 2 | Ferret<br>(*Mustela putorius furo*) | 2 | Paul-Ehrlich-Institut, Langen, Germany | 12.5 ng/ml | 100 nM | 0.48 µg/ml | 2.5 µM |
| 3 | Cat<br>(*Felis catus*) | 2 | Justus-Liebig-University, Giessen, Germany | 25 ng/ml | 50 nM | 0.072 µg/ml | - |
| 4 | Dog<br>(*Canis lupus familiaris*) | 1 | Institute of Animal Pathology, Bern, Switzerland | 25 ng/ml | 50 nM | 0.072 µg/ml | - |
| 5 | Rabbit<br>(*Oryctolagus cuniculus*) | 1 | Slaughterhouse, Bern, Switzerland | 25 ng/ml | 50 nM | 0.48 µg/ml | - |
| 6 | Pig<br>(*Sus scrofa domestica*) | 2 | Institute of Virology and Immunology, Mithelhäusern, Switzerland | 25 ng/ml | 70 nM | 0.072 µg/ml | - |
| 7 | Cattle (*Bos Taurus*) | 1 | Institute of Animal Pathology, University of Bern, Switzerland | 25 ng/ml | 50 nM | 0.48 µg/ml | 2.5 µM |
| 8 | Goat (*Capra aegagrus hircus*) | 2 | Slaughterhouse, Bern, Switzerland | 12.5 ng/ml | 50 nM | 0.48 µg/ml | 2.5 µM |
| 9 | Bactrian camel<br>(*Camelus bactrianus*) | 1 | Institute of Animal Pathology, University of Bern, Switzerland | 5 ng/ml | 50 nM | 0.072 µg/ml | - |
| 10 | Llama (*Llama glama*) | 2 | Institute of Animal Pathology, University of Bern, Switzerland | 5 ng/ml | 50 nM | 0.072 µg/ml | - |
| 11 | Bat (*Sturnira lilium*) | 1 | Costa Rica, (CIET-315-2013; Permit 1841/14) | 5 ng/ml | 50 nM | 0.48 µg/ml | - |
| 12 | Bat (*Carollia perspicillata*) | 1 | Costa Rica (CIET-315-2013; Permit 1841/14) | 5 ng/ml | 50 nM | 0.48 µg/ml | - |

The quantification of the viral RNA load at both temperatures showed that there was a progressive 4-log fold increase in viral RNA load at 72- and 96-hours post-infection (hpi) in rhesus macaque and cat AEC cultures. In contrast, for the remaining animal AEC cultures either a continuous or declining level of viral RNA load was detected throughout the entire time course (**Fig. 1a and b**, **Fig. S1b and c**). Because molecular assays do not discern between infectious and non-infectious virus, we also performed viral titration assays with the corresponding apical washes ^21^. This corroborated our previous findings, namely that only AEC cultures derived from rhesus macaque and cat displayed a progressive increase in viral SARS-CoV-2 titers over time, while for the majority of species no infectious virus was detected beyond 24 hpi **(Fig. 1c and d**, **Fig. S1d and e)**. The viral titers observed in the rhesus macaque and cat AEC cultures are comparable to those that we previously observed for human AEC cultures, where we also observed a 4-log fold rise of progeny released virus in the apical side ^20^. Interestingly, although ferrets have previously been shown to be susceptible to SARS-CoV-2, we observed no viral replication in AEC cultures derived from the tracheobronchial regions. Instead, only low levels of SARS-CoV-2 viral titers at 72 and 96 hpi were detected at 37°C, in agreement with *in vivo* studies in ferrets showing a dose-dependent and limited SARS-CoV-2 infection restricted to the upper respiratory tract 22–24. We further analysed SARS-CoV-2 infection in the animal AEC cultures by staining for SARS-CoV-2 nucleocapsid protein on formalin-fixed AEC cultures, to visualize intracellular presence of the virus. This revealed that SARS-CoV-2 antigen-positive cells were detected in rhesus macaque and cat AEC cultures at 96 hpi, while in the other animal AEC cultures, including those of ferrets, no SARS-CoV-2 antigen-positive cells were observed (**Fig. 2**, **Fig. S1a**). This further confirmed that amongst animals studied, only monkey and cat airway epithelial cells support efficient replication of SARS-CoV-2.

**Figure 1.**
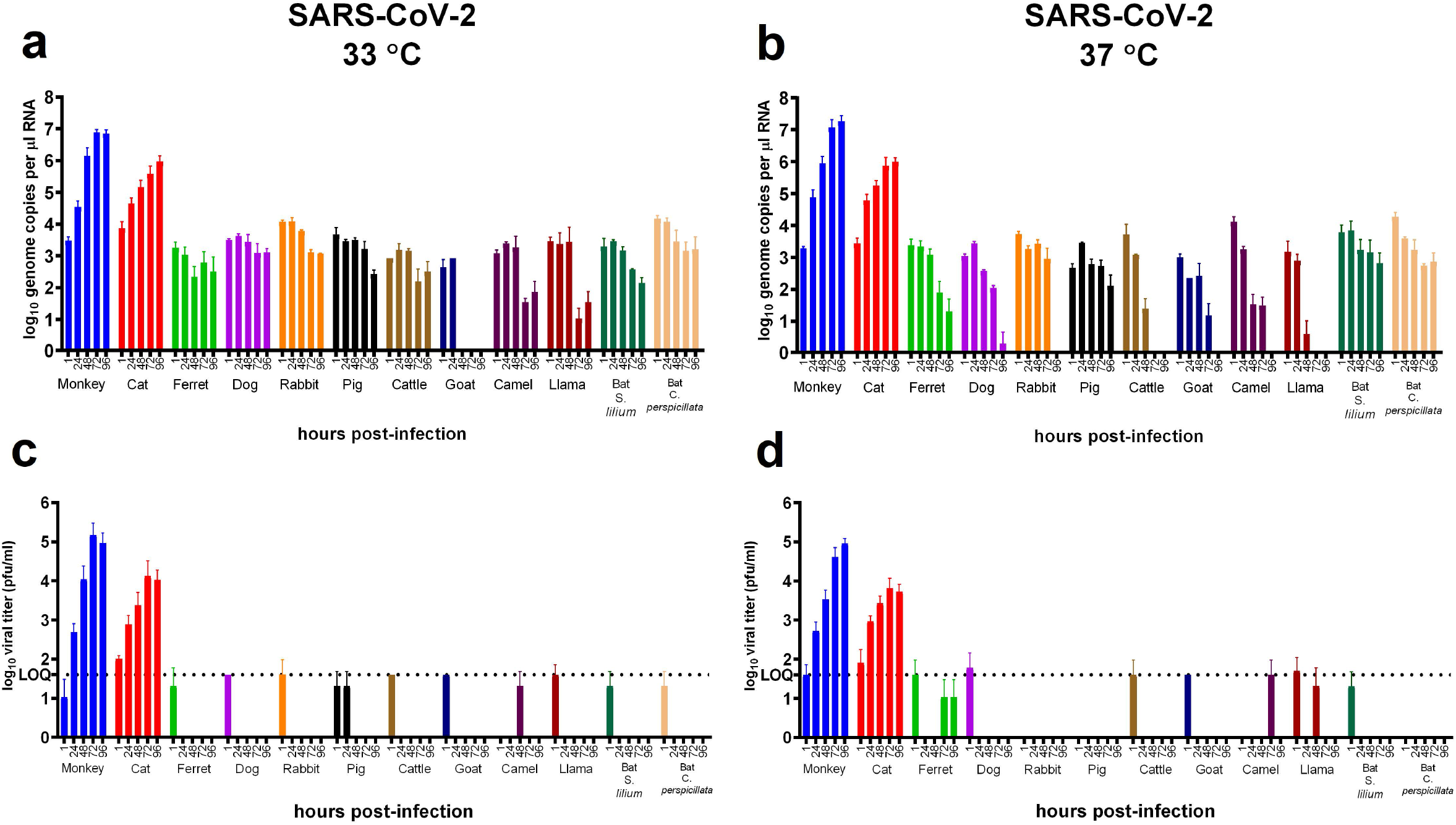
SARS-CoV-2 replication kinetics in diverse mammalian species. Well-differentiated animal AEC cultures derived from the tracheobronchial epithelial cells were inoculated with 30.000 PFU of SARS-CoV-2 at either 33°C or 37°C. Inoculated virus was removed at 1 hpi and the apical side was washed three times. Cultures were further incubated for 96 h. At the indicated time post infection, apical virus release was assessed by qRT-PCR targeting the E gene (**a, b**) and plaque titration assays on Vero E6 cells (**c, d**). Error bars represent the average of two independent biological replicates using AEC cultures established from one or two biological donors. The dotted lines on figure **c** and **d** indicate the detection limit of the assay.

**Figure 2.**
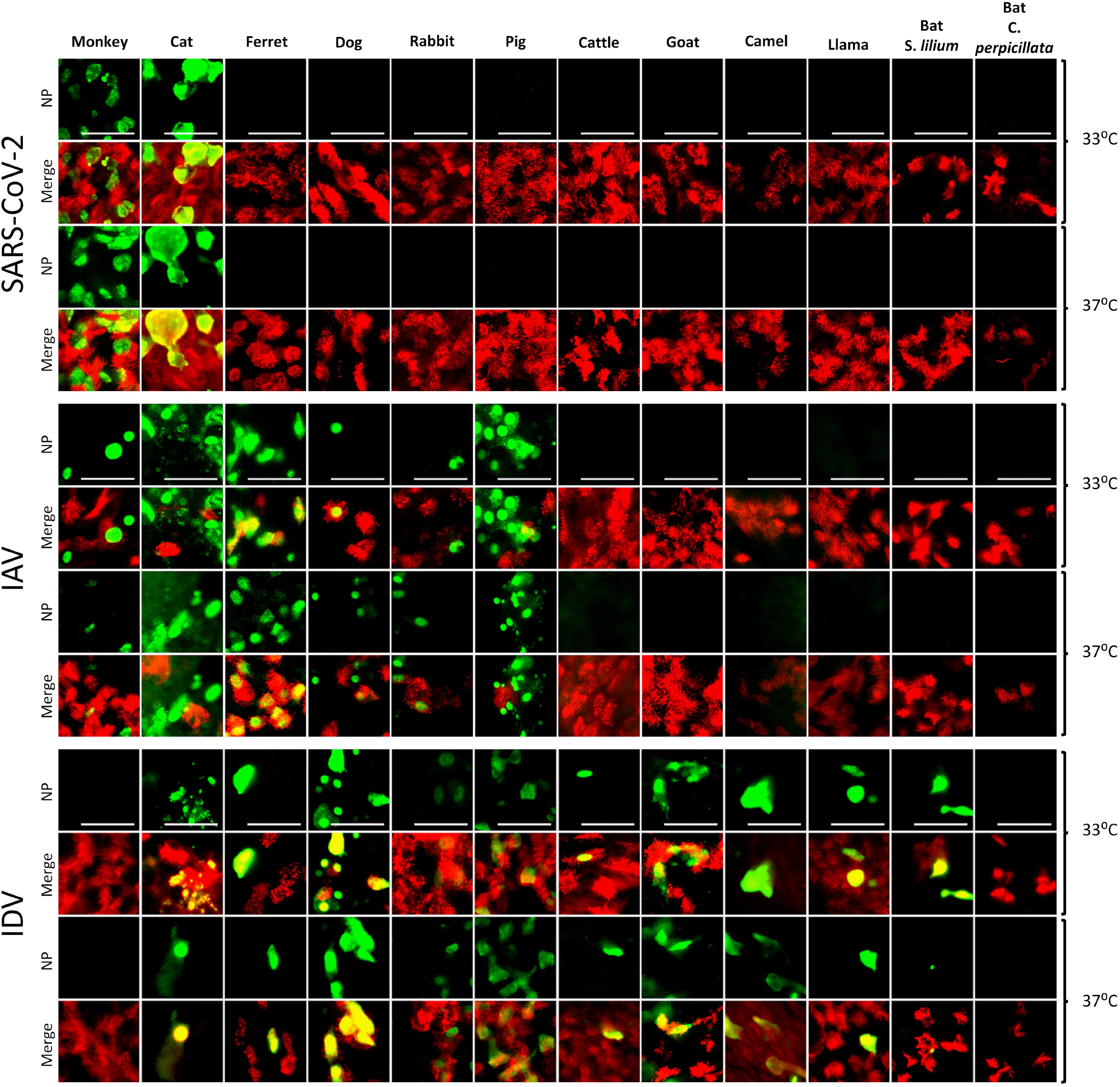
Tropisms of SARS-CoV-2, IAV, and IDV in infected AEC cultures from diverse mammalian species. Well-differentiated animal AEC cultures were inoculated with either 30.000 pfu of SARS-CoV-2 (SARS-CoV-2/München-1.1/2020/929), 10.000 TCID_50_ of IAV/Hamburg/4/2009 (H1N1pdm09) or IDV (D/bovine/Oklahoma/660/2013). Virus-infected AEC cultures were incubated at respectively 33°C or 37°C and fixed 96 hpi (for SARS-CoV-2) or 48 hpi (for IAV and IDV). Following fixation, virus-infected cultures were stained using antibodies against either SARS-CoV-2, IAV, or IDV Nucleocapsid protein (green), and β-tubulin (cilia, red). Images were acquired using an EVOS FL Auto 2 Imaging System equipped with a 40x air objective. Scale bar = 50 μm.

The absence of infectious progeny virus in most animal species, except rhesus macaques and cats, indicates that certain animal species may be intrinsically refractory to SARS-CoV-2 infection, which may be due to incompatibility with the cellular receptor utilized by SARS-CoV-2 for cellular entry 25,26. To assess whether the observed susceptibility to SARS-CoV-2 corresponds to the amino acid sequence conservation of the receptor-binding motif (RBM) in ACE2 we performed *in silico* analysis on the available ACE2 protein sequences ^25,27^. The ACE2 protein sequences from the two neotropical bat species (*S. lilium* and *C. perspicillata*) were not included in the analysis, due to their unavailability. Similarly, the ACE2 protein sequence for llama is not available, and therefore we used the sequence of alpaca (XM_006212647.3) as an alternative, as it is the closest relative. This revealed that in comparison to humans, the ACE2 RBM regions interacting with the receptor-binding domain (RBD) of SARS-CoV-2 are well conserved in rhesus macaques and cats while being slightly more diverse in other species (**Fig. S2a**).

Apart from receptor compatibility as a limiting factor of virus infection, it has been demonstrated previously that partially differentiated AEC cultures are poorly permissive to respiratory virus infection ^28^. To investigate whether the lack of replication in for instance the ferret cells, was not caused by poor differentiation of our cell cultures, we validated the AEC cultures by performing infections with the 2009 pandemic influenza A virus (A/Hamburg/4/2009, IAV) and the ruminant-associated influenza D virus ((D/bovine/Oklahoma/660/2013, IDV). Both viruses are members of *Orthomyxoviridae* and are known to have a broad host spectrum, including ferrets ^15–17,29^. The AEC cultures from 12 different species (rhesus macaque, cat, ferret, dog, rabbit, pig, cattle, goat, llama, camel, and two neotropical bats) were inoculated with 10.000 TCID_50_ of either IAV or IDV and incubated at 33°C and 37°C. After 48 hours, the AEC cultures were fixed and processed by immunofluorescence assays. This analysis showed that, in contrast to SARS-CoV-2, IAV antigen-positive cells could be detected in both companion animals AEC cultures, as well as in the commonly used animal models, such as ferret, monkey, rabbit, and porcine AEC cultures **(Fig. 2**, **Fig. S1a)**^30^. For IDV we observed antigen-positive cells in all AEC model, except for rhesus macaque and one of the neotropical bat species, indicating that the AEC cultures were all well-differentiated and susceptible to virus infection.

In the immunofluorescence analysis we also incorporated an antibody against beta-tubulin marker to discern ciliated and non-ciliated cell populations. For both rhesus macaques and cats, SARS-CoV-2 antigen-positive cells predominantly overlapped with the non-ciliated cell populations, irrespective of the incubation temperature. Using a polyclonal antibody against the human ACE2 we could observe that the cellular receptor expression in rhesus macaques predominantly overlaps with SARS-CoV-2 cell tropism, indicating same cellular tropism (**Fig. S2b**). Unfortunately, the polyclonal antibody against the human ACE2 did not bind the feline ACE2 protein, we could therefore not formally demonstrate that SARS-CoV-2 virus-infected cat cells are indeed expressing ACE2 on their surface.

### Whole-genome sequence analysis

It has previously been shown that SARS-CoV-2 can undergo rapid genetic changes *in vitro* ^31^. Since we observed efficient replication in rhesus macaque and cat AEC cultures, therefore we assessed whether any mutations suggestive of viral adaptation had occurred. We performed whole-genome sequencing (Nanopore sequencing technology) on the viral inoculum used as well as the progeny viruses collected after one passage, at 96 hpi from the rhesus macaque and cat AEC cultures incubated at 33°C and 37°C. This inoculum was either passage 1 or passage 2 virus stocks from the SARS-CoV-2/München-1.1/2020/929 isolate we had received. In the viral sequences in the 96 hpi samples from virus-infected rhesus macaque and cat AEC cultures, we observed no obvious signs of nucleotide transitions that lead to nonsynonymous mutations compared to the respective inoculums (**Fig. 3**), irrespective of temperature and animal species. This highlights that the currently circulating SARS-CoV-2 D614G-variant can productively infect rhesus macaque and cat airway epithelial cells.

**Figure 3.**
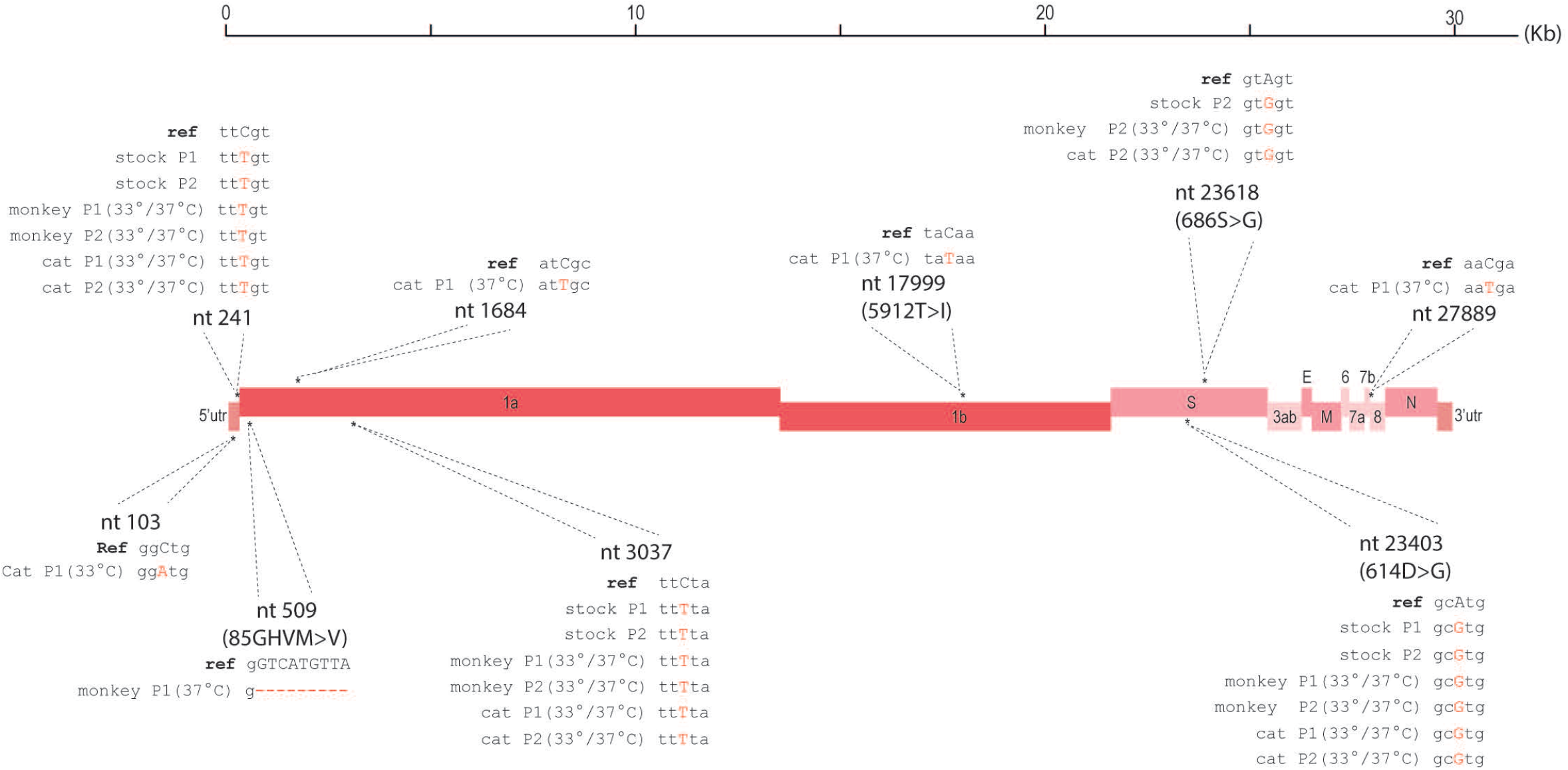
Whole-genome sequencing analysis using Nanopore sequencing technology. A graphical representation of variants found in the SARS-CoV-2 stock passage 1 (P1) and 2 (P2), as well as the apical washes from SARS-CoV-2-infected monkey and cat AEC cultures with either P1 or P2 stock 96 hpi at 33°C or 37°C. SARS-CoV-2/Wuhan-Hu-1 (MN908947.3) was used as the reference sequence.

## Discussion

This is the first study employing an *in vitro* AEC culture repository composed of various domestic and wildlife animal species to assess the potential intermediate and spillback host reservoir spectrum of SARS-CoV-2. Inoculation of AEC cultures of rhesus macaque, cat, ferret, dog, rabbit, pig, cattle, goat, llama, camel, and two neotropical bat species with SARS-CoV-2 revealed that only tracheobronchial cells from rhesus macaque and cats supported efficient replication of SARS-CoV-2. Whole-genome sequencing indicated that the current circulating SARS-CoV-2 D614G-variant can efficiently infect rhesus macaque and cat airway epithelial cells. Our data highlight that these two animals are potential models for the evaluation of therapeutic mitigation strategies for currently circulating viral variants. In conjunction with the previous documented spillover events, close surveillance of these animals, including closely related species, in the wild, captivity, and household situations is warranted.

To date, there have been several published reports evaluating the suitability of animal models towards SARS-CoV-2 infection, including cats, rhesus macaques, dogs, pigs, and ferrets ^22,23,32–34^. Interestingly, we observed that SARS-CoV-2 does not efficiently replicate in our tracheobronchial airway epithelial cells derived from ferrets, whereas ferrets are used as animal models. This may be due to viral infections in ferrets are mainly restricted to the nasal conchae and are dose-dependent, and additionally, the origin of the cells used as input for the AEC may not recapitulate the cells of the nasal mucosa ^24,32,34^. It is known that there are differences in cellular composition and the host determinant expression levels along proximal and distal regions of the respiratory tract ^35^. Additionally, SARS-CoV-2 may even utilize a different cellular receptor in ferrets ^36^. Therefore, it would be of interest to complement our current repository with AEC cultures from different anatomical regions of animals like ferrets and to evaluate whether ACE2 is employed by SARS-CoV-2 as the cellular receptor in the different animal species.

It has been proposed that SARS-CoV-2 spillover into the human population, like SARS-CoV, has originated from bats, either directly or via an intermediate reservoir ^3,37^. With more than 1200 bat species comprising more than 20% of all mammalian species, we restricted our experiments with SARS-CoV-2 to our established AEC cultures from the two neotropical *Carollia perspicillata* and *Sturnira lilium* bat species (*Gultom et al Manuscript in preparation*). We show that these two neotropical bats are not susceptible to SARS-CoV-2, suggesting that they are not a likely reservoir host for SARS-CoV-2, despite the detection of other coronaviruses and presumptive ACE2 receptor usage by SARS-CoV-2 in the closely related bat species ^38,39^. Interestingly, it has recently been described that fruit bats (*Rousettus aegyptiacus*) are susceptible to SARS-CoV-2 infection ^23^. Future research should therefore include AEC cultures from these bats, as well as from horseshoe bat species (genus Rhinolophus), as they have previously been characterized as a reservoir host for viruses that have a close genetic relationship with the coronavirus associated with the SARS outbreak in 2002/2003 ^23,37^.

Taken together, our results highlight that *in vitro* well-differentiated airway epithelial models in combination with high throughput genomic analysis can be applied as viable surrogate models to refine, reduce, and replace animal experimentation to evaluate host tropism of respiratory viruses, and thereby providing important insight into the host spectrum of SARS-CoV-2.

## Materials and Methods

### Conventional cell culture

Vero E6 cells (kindly provided by Doreen Muth, Marcel Müller, and Christian Drosten, Charité, Berlin, Germany) were cultured in Dulbecco’s Modified Eagle Medium (DMEM) supplemented with 10% (v/v) heat-inactivated fetal bovine serum (FBS), 1 mM sodium pyruvate, 1x GlutaMAX, 100 μg/ml streptomycin, 100 IU/ml penicillin, 1% (v/v) non-essential amino acids, and 15 mM HEPES (Gibco). Cells were maintained at 37°C in a humidified incubator with 5% CO_2_.

### Establishment of animal airway epithelial cell (AEC) culture repository

Tracheobronchial epithelial cells from 12 different animal species were isolated from post-mortem tracheobronchial tissue that was obtained in collaboration with slaughterhouses, veterinary hospitals, and both domestic and international research institutes that euthanize their animals for diagnostic purposes, or as part of their licensed experimental work in accordance with local regulations and ethical guidelines. Isolation and culturing was performed as previously described ^18^. For the establishment of well-differentiated AEC cultures from diverse mammalian species, several modifications to the composition of the ALI medium were introduced (**Table. 1**). All animal ALI cultures were maintained at 37°C in a humidified incubator with 5% CO_2_. During the development of differentiated ALI cultures (3-4 weeks), media was changed every 2-3 days.

### Viruses

SARS-CoV-2 (SARS-CoV-2/München-1.1/2020/929) was kindly provided by Marcel Müller and Christian Drosten. The virus stock was propagated in Vero E6 for 48 hours. Virus containing supernatant was cleared from cell debris through centrifugation for 5 minutes at 500 × g before aliquoting and storage at −80°C. The viral titer was determined by Plaque Forming Unit (PFU) assay on Vero E6 as previously described ^20,40^.

The Influenza A/Hamburg/4/2009 (H1N1pdm09) virus strain in the pHW2000 reverse genetic backbone was kindly provided by Martin Schwemmle, University of Freiburg, Germany. Working stocks were prepared by propagating the rescued virus in MDCK-II cells for 72 hours in the infection medium, which is composed of Eagle’s Minimum Essential Medium (EMEM), supplemented with 0,5% of BSA (Sigma-Aldrich), 100 μg/ml Streptomycin and 100 IU/ml Penicillin (Gibco), 1 μg/ml Bovine pancreas-isolated acetylated trypsin (Sigma-Aldrich), and 15 mM HEPES. The viral titer was determined by TCID_50_ assay on MDCK-II cells as described previously ^41,42^.

Influenza D virus (D/bovine/Oklahoma/660/2013) was kindly provided by Feng Li, South Dakota University, United States. Virus stocks were propagated in the human rectal tumor cell line HRT-18G (CRL11663, ATCC) for 96 hours in the infection medium, with the adjustment of using 0,25 μg/ml of trypsin. The viral titer was determined by TCID_50_ assay on HRT-18G cells as previously described ^43^.

### Infection of animal AEC cultures

Well-differentiated AEC cultures from 12 different species were infected with 30.000 PFU of SARS-CoV-2, or 10.000 TCID_50_ of either IAV or IDV. Viruses were diluted in Hanks balanced salt solution (HBSS, Gibco), inoculated via the apical side, and incubated for 1 h, at either 33°C or 37°C. Afterwards, inoculum was removed, and the apical surfaces were rinsed three times with HBSS. Virus-infected and control AEC cultures were incubated at the indicated temperatures in a humidified incubator with 5% CO_2_. Progeny virus release was monitored with 24-hour intervals for a total duration of 96 hours, through the application of 100 μl of HBSS onto the apical surface and incubated 10 min prior to the collection time point. The collected apical washes were diluted 1:1 with virus transport medium (VTM) and stored at −80°C for later analysis. Following the collection of the apical washes, the basolateral medium was exchanged with fresh ALI medium. Each experiment was repeated as two independent biological replicates using AEC cultures established from either one or two biological donors of each species, depending on the availability of procured animal tissue (**Table. 1**).

### Immunofluorescence analysis

Virus-infected animal AEC cultures were fixed with 4% (v/v) neutral-buffered formalin at 96 hpi for SARS-CoV-2 or 48 hpi for IAV/IDV infected AEC cultures and processed as previously described ^18^. For the detection of SARS-CoV-2, fixed animal AEC cultures were incubated with a rabbit polyclonal antibody against SARS-CoV Nucleocapsid protein (Rockland, 200-401-A50), which has previously been shown to cross-react with SARS-CoV-2 ^20^. To detect the presence of IAV and IDV virus-infected cells, a mouse antibody against Influenza A Virus NP Protein (clone C43; ab128193, Abcam) and a custom-made rabbit polyclonal antibody against the NP of influenza D/bovine/Oklahoma/660/2013 strain (Genscript, Piscataway, NJ, USA) were used, respectively. To visualize the distribution of ACE2 in the AEC cultures, a rabbit polyclonal antibody against ACE2 (ab15348, Abcam) was used. Alexa Fluor® 488-labeled donkey anti-Rabbit and -mouse IgG (H + L) (Jackson Immunoresearch) were used as secondary antibodies. Alexa Fluor® 647-labeled rabbit anti-β-tubulin (9F3, Cell Signaling Technology and EPR16775, Abcam) and Alexa Fluor® 594-labeled mouse anti-ZO1 (1A12, Thermo Fisher Scientific) were used to visualize cilia and tight junctions, respectively. All samples were counterstained using 4’,6-diamidino-2-phenylindole (DAPI, Thermo Fisher Scientific) to visualize the nuclei. Imaging was performed using EVOS FL Auto 2 Imaging System equipped with a Plan Apochromat 40x/0.95 air objective. All images were processed using Fiji software packages ^44^. Brightness and contrast were adjusted identically to their corresponding controls. Figures were assembled using the FigureJ plugin ^45^.

### Quantitative Real-Time Reverse Transcription Polymerase Chain Reaction (qRT-PCR)

Viral RNA was extracted from 100 μL of 1:1 diluted apical wash using the NucleoMag VET (Macherey-Nagel AG, Oensingen, Switzerland), according to the manufacturer’s guidelines, on a Kingfisher Flex Purification system (Thermo Fisher Scientific, Darmstadt, Germany). Two microliters of extracted RNA were amplified using TaqMan™ Fast Virus 1-Step Master Mix (Thermo Fisher Scientific) according to the manufacturer’s protocol. For the detection we used a forward primer 5′-ACAGGTACGTTAATAGTTAATAGCGTACTTCT-3′, reverse 5′- ACAATATTGCAGCAGTACGCACA −3′ and probe 5′-FAM-ATCCTTACTGCGCTTCGA-MGB-Q530-3′ (Microsynth, Balgach, Switzerland) targeting the Envelope gene of SARS-CoV (AY291315.1) and SARS-CoV-2 (MN908947.3), which is an adapted version of primers described by Corman and colleagues ^46^. As a positive control, a serial dilution of *in vitro* transcribed RNA containing regions of the RdRp, E and N gene, derived from a SARS-CoV-2 synthetic construct (MT108784) were included to determine the genome copy number ^40^. Measurements and analysis were performed using an ABI7500 instrument and associated software package (Applied Biosystems, Foster City, CA, USA).

### Titration of SARS-CoV-2 in the apical washes

For the quantification of SARS-CoV-2 apical washes were titrated by plaque assay on Vero E6 cells. Briefly, 1×10^5^ cells/well were seeded in 24-well plates one day prior to the titration and inoculated with 10–fold serial dilutions of virus solutions. Inoculums were removed 1 hpi and replaced with overlay medium consisting of DMEM supplemented with 1.2% Avicel (RC-581, FMC biopolymer), 10% heat-inactivated FBS, 100 μg/ml streptomycin, and 100 IU/ml penicillin. Cells were incubated at 37 °C with 5% CO_2_ for 48 hours and fixed with 4% (v/v) neutral buffered formalin prior to staining with crystal violet ^47^.

### ACE2 homology analysis

For the analysis on the conservation of ACE2 among different species, the available ACE2 protein sequences from human (NM_001371415.1), rhesus macaque (NM_001135696.1), cat (XM_023248796.1), ferret (NM_001310190.1), dog (NM_001165260.1), rabbit (XM_002719845.3), pig (NM_001123070.1), cattle (XM_005228428.4), goat (NM_001290107.1), Bactrian camel (XM_010968001.1), and alpaca (XM_006212647.3). The protein alignment was performed using ClustalW in Geneious 11.1.5. Software (Biomatters) using the default setting. ACE2 protein residues interacting with SARS-CoV-2, receptor binding motif (RBM), were selected based on previous described critical ACE2 residues interacting with SARS-CoV-2 receptor binding domain (RBD) ^25,27^.

### Whole-Genome Sequencing using Oxford Nanopore (MinION)

Sequencing was performed on viral RNA isolated from the SARS-CoV-2 stock and the 96 hpi apical washes of SARS-CoV-2-infected monkey and cat AEC cultures according to the ARTIC platform nCoV19 protocols ^48,49^. The v2 protocol was used as a basis for the reverse transcript and tiled multiplex PCR reaction using the ARTIC nCoV-2019 V3 primer pool (see **Table. S1**), whereas the v3 protocol was used for the downstream library preparation. Sequencing libraries were generated using the Native Barcoding Expansion 96 kit (EXP-NBD196, Oxford Nanopore Technologies) and sequenced on a MinION R9.4.1 flowcell according to the manufacturer's instructions for a duration of 48 hours. Data acquisition and real time high accuracy base-calling was performed using MinION software (v20.06.4). Demultiplexing and read filtering were done according to the ARTIC platform nCoV19 pipeline and the consensus calling was performed using the experimental Medaka pipeline. Consensus sequences were aligned and further analyzed in Geneious 11.1.5. (Biomatters) using SARS-CoV-2/Wuhan-Hu-1 (MN908947.3) as the reference sequence.

## Acknowledgements

We gratefully thank Lia van der Hoek, University Medical Center, Amsterdam, the Netherlands, for critical reading of the manuscript and helpful discussions. This study was supported by research grants from the European Commission (Marie Sklodowska-Curie Innovative Training Network “HONOURS”; grant agreement No 721367), the Swiss National Science Foundation (SNF grants 310030_179260 and 31CA30_196062), the Federal Food Safety and Veterinary Office (FSVO grant 1.20.02), and the German Research Foundation (DFG, SFB 1021, project B01).

## Author contributions

Conceptualization, R.D; Investigation, M.G, M.L, L.L, M.W, M.S, S.S, A.K, HP.S, J.P, M.H, P.V, and T.T; Resources, N.E, N.SR, C.G, P.Z, H.P, S.SC, G.T, N.R, A.VS, A.MS, E.CA, V.vM, and B.S,; Writing, M.G, R.D; Reviewing and Editing, R.D; Visualization; M.G, M.L, T.T; Supervision, V.T and R.D; Funding Acquisition, V.T and R.D.

---

**Figure S1. Mock-treated cells from the infection of animal AEC cultures with SARS-CoV-2, IAV, and IDV.** Mock treated AEC cultures were incubated at 33°C or 37°C in parallel with virus-infected cells for 96 hpi (for SARS-CoV-2) or 48 hpi (for IAV and IDV). Afterwards, cells were fixed and stained using antibodies against either SARS-CoV, IAV, or IDV Nucleocapsid protein (green), and β-tubulin (cilia, red), and tight-junctions (white) **(a)**. Images were acquired using an EVOS FL Auto 2 Imaging System equipped with a 40x air objective. Scale bar = 50 μm. In parallel with the SARS-CoV-2-infected cells, apical washes from the mock-treated cells were collected every 24h and analysed by qRT-PCR (**b, c**) and plaque titration assays on Vero E6 cells (**d, e**). Error bars represent the average of two independent biological replicates using AEC cultures established from one or two biological donors. The dotted lines on figure **d** and **e** indicate the detection limit of the assay.

**Figure S2. ACE2 analysis amongst different animal species.** Protein sequence alignment of ACE2 from diverse animals in the residues interacting with SARS-CoV2 **(a)**. The alignment was constructed using ClustalW program. To visualize the ACE2 distribution in monkey AEC cultures, formalin-fixed cells were stained with antibodies against ACE2 (green), β-tubulin (cilia, red), and ZO-1 (tight junctions, white) **(b)**. Image acquisition was performed using an EVOS FL Auto 2 Imaging System equipped with a 40x air objective. Scale bar = 50 μm.

