## Supplementary figures and images for "Susceptibility of well-differentiated airway epithelial cell cultures from domestic and wildlife animals to SARS-CoV-2"

### Figure S1

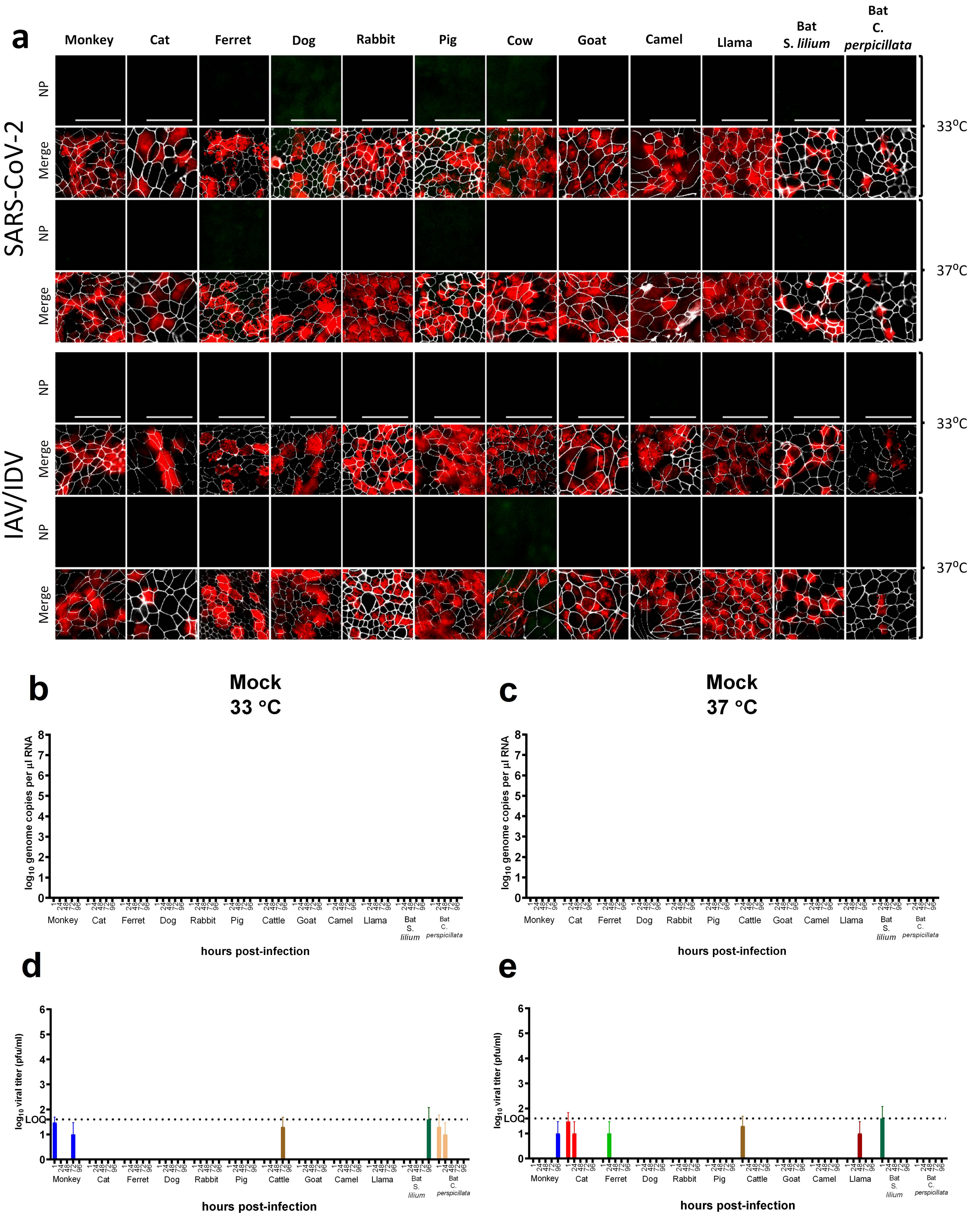
