## Supplementary material for "Susceptibility of well-differentiated airway epithelial cell cultures from domestic and wildlife animals to SARS-CoV-2": Figure S2

**A**

| | 19 | 24 | 27 | 28 | 30 | 31 | 34 | 35 | 37 | 38 | 41 | 42 | 45 | 49 | 79 | 82 | 83 | 90 | 322 | 325 | 329 | 330 | 353 | 354 | 355 | 357 | 393 |
|---|---|---|---|---|---|---|---|---|---|---|---|---|---|---|---|---|---|---|---|---|---|---|---|---|---|---|---|
| Human (Homo sapiens) | S | Q | T | F | D | K | H | E | E | D | Y | Q | L | N | L | M | Y | N | N | Q | E | N | K | G | D | R | R |
| Monkey (Rhesus macaque) | · | · | · | · | · | · | · | · | · | · | · | · | · | · | · | · | · | · | · | · | · | · | · | · | · | · | · |
| Cat (Felis catus) | · | L | · | · | E | · | · | · | · | E | · | · | · | · | · | T | · | · | · | · | · | · | · | · | · | · | · |
| Ferret (Mustela putorius furo) | · | L | · | · | E | · | Y | · | · | E | · | · | · | · | H | T | · | D | · | E | Q | · | · | R | · | · | · |
| Dog (Canis lupus familiaris) | · | L | · | · | E | · | Y | · | · | E | · | · | · | · | · | T | · | D | · | · | G | · | · | · | · | · | · |
| Rabbit (Oryctolagus cuniculus) | · | L | · | · | E | · | Q | · | · | · | · | · | · | D | · | T | · | · | S | · | · | · | · | · | · | · | · |
| Pig (Sus scrofa) | · | L | · | · | E | · | L | · | · | · | · | · | · | T | I | T | · | T | · | · | N | · | · | · | · | · | · |
| Cow (Bos taurus) | · | · | · | · | E | · | · | · | · | · | · | · | · | · | M | T | · | · | Y | · | D | · | · | · | · | · | · |
| Goat (Capra hircus) | · | · | · | · | E | · | · | · | · | · | · | · | · | · | M | T | · | · | Y | · | N | · | · | · | · | · | · |
| Camel (Camelus bactrianus) | · | L | · | · | E | E | · | · | · | · | · | · | · | · | T | T | · | · | · | · | D | · | · | · | · | · | · |
| Alpaca (Vicugna pacos) | · | L | · | · | K | E | · | · | · | · | · | · | · | · | A | I | · | · | · | · | D | · | · | · | · | · | · |

**B**

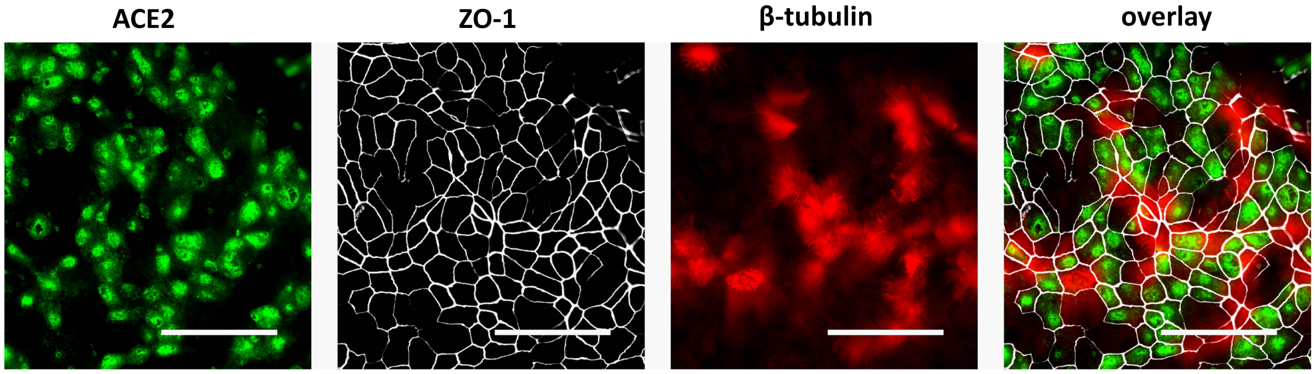
